# Phylogenetic diversity of light dependent phosphorylation of Thr78 in Rca

**DOI:** 10.1101/2024.08.27.609719

**Authors:** Nikita Bhatnagar, Sarah S. Chung, John Hodge, Ryan A Boyd, Sang Yeol Kim, Mia Sands, Andrew D. B. Leakey, Donald R. Ort, Steven Burgess

## Abstract

Rubisco activase is an ATP-dependent chaperone that facilitates dissociation of inhibitory sugar phosphates from the catalytic sites of ribulose-1,5-bisphosphate carboxylase/oxygenase during photosynthesis. In Arabidopsis, Rubisco activase is negatively regulated by dark-dependent phosphorylation of threonine 78. The prevalence of threonine 78 in Rubisco activase was investigated across sequences from 91 plant species, finding 29 (∼32%) species shared a threonine in the same position. Analysis of seven C3 species with an antibody raised against a threonine 78 phospho-peptide demonstrated that this position is phosphorylated in multiple genera. However, light-dependent dephosphorylation of threonine 78 was observed only in Arabidopsis. Further, phosphorylation of threonine 78 could not be detected in any of the four C4 grass species examined. The results suggest that despite conservation of threonine 78 in Rubisco activase from a wide range of species, a regulatory role for phosphorylation at this site is more limited. This provides a case study for how variation in post-translational regulation can amplify functional divergence across the phylogeny of plants beyond what is explained by sequence variation in a metabolically important protein.

## INTRODUCTION

Rubisco catalyzes the addition of CO_2_ to a molecule of ribulose-1,5-bisphosphate (RuBP) forming two molecules of 3-phosphoglycerate (3-PGA) and in many circumstances is the rate limiting step of CO_2_ assimilation (Stitt *et al*., 1991; Hudson *et al*., 1992; Stitt & Schulze, 1994; Poolman *et al*., 2000; Zhu *et al*., 2007). To perform this reaction the enzyme must first be activated through attachment of CO_2_ to a catalytic lysine in order to bind Mg^2+^ (Lorimer and Miziorko, 1980). However, the enzyme is susceptible to inhibition by attachment of RuBP prior to activation, or binding of one of several inhibitory sugar phosphates to the active site (Orr et al., 2023). Inhibitors can be released by the action of Rubisco activase (Rca) (Somerville et al., 1982; Salvucci et al., 1986; Jiang et al., 1994), which uses energy from ATP hydrolysis to drive conformational changes in Rubisco, freeing the active site (Neuwald et al., 1999; Portis & Jr., 2003; Flecken et al., 2020). Therefore, Rca is critical for regulation of Rubisco activity and carbon assimilation (Parry et al., 2013; Wijewardene et al., 2021; Qu et al., 2023; Orr et al., 2023).

Rca itself is subject to regulation, which provides a link between carbon assimilation and electron transport. Most of our understanding about this process comes from Arabidopsis, where there are two isoforms of Rca which differ in size. The longer ‘α’ isoform possesses a 20-30 amino acid C-terminal extension (To et al., 1999) which contains two cysteine residues (Cys-451 and Cys-470). The cysteines are regulated by thioredoxin f in response to changes in the ATP/ADP arising from variation in rates of electron transport and ATP synthesis (Zhang & Portis, 1999b; Zhang et al., 2002). In the dark, the cysteines are oxidized to form a disulfide bridge (oxidized state, E_m ∼_344 mV pH 7.9) (Hutchison et al., 2000), increasing the sensitivity of Rca to inhibition by ADP (Portis et al., 2007), downregulating Rca activity and CO_2_ fixation. However, redox regulation of Rca does not appear to be essential. Solanaceae species, such as tobacco, are reported to only express Rca-β, which lacks the C-terminal extension, (Nagarajan & Gill, 2018), and the sensitivity of Rca-β isoforms to inhibition by ADP varies (Carmo-Silva & Salvucci, 2013). Therefore, it is apparent that additional regulatory mechanisms are also important to control Rca activity.

Rca a and β isoforms, which differ in ADP sensitivity, can be produced by both alternate splicing (Werneke et al., 1989; Rundle & Zielinski, 1991; To et al., 1999) or from transcription of independent genes (Rundle & Zielinski, 1991; Salvucci et al., 2003). In addition, the relative expression level and catalytic properties of Rca-α and Rca-β differ across species (Rundle & Zielinski, 1991; Crafts-Brandner et al., 1997; Carmo-Silva & Salvucci, 2013; Shivhare & Mueller-Cajar, 2017; Degen et al., 2021) and can be impacted by environmental factors such as temperature (Shivhare & Mueller-Cajar, 2017; Scafaro et al., 2019; Kim et al., 2020, 2021; Degen et al., 2021) to modulate CO_2_ fixation under variable conditions.

Post-translational modification of Rca has also been proposed to provide an additional layer of regulation (Amaral et al., 2024). In Arabidopsis, Threonine 78 (Thr78) is phosphorylated in the dark and under low light intensities, but not under high light, suggesting a regulatory role (Reiland *et al*., 2009; Boex□Fontvieille *et al*., 2014; Kim *et al*., 2016). Functional support for this hypothesis came from several findings: (1) phosphorylation at Thr78 (Thr78-Rca) in Arabidopsis is reported to cause a significant decrease in plant biomass, due to a decreased Rubisco activation state (Kim *et al*., 2019); (2) transgenic Arabidopsis plants expressing phospho-null mutants of Rca (T78A) had enhanced photosynthetic efficiency and plant growth; and (3) substitution of Thr78-to-Ser (T78S) resulted in hyper-phosphorylation, leading to reduced carbon assimilation and slower growth (Kim *et al*., 2019). cpCK2 was shown to be the major kinase responsible for dark phosphorylation of AtRca, with loss-of-function mutant lines showing a reduction in Rca phosphorylation (Kim *et al*., 2016). However, it is not clear whether this proposed mechanism is relevant outside of Arabidopsis. To address this issues, we sought to define the prevalence of this post-translational modification in photosynthetic organisms. This was achieved by considering the following four questions, (1) at what stage of flowering plant evolution did Thr78 emerge? (2) how conserved is Thr78 among photosynthetic angiosperms? (3) to what extent is Thr78 phosphorylated in different species? (4) and is light-dependent de-phosphorylation of Thr78 a general mechanism of controlling of RCA activity?

## MATERIAL AND METHODS

### Determination of Rca isoform and photosynthetic species containing Thr78-Rca

To analyze the conservation of Thr78 in Rca protein across plant species, the sequence of Arabidopsis Rca (AtRca: AT2G39730) was retrieved from the *Arabidopsis thaliana* TAIR10 genome assembly (Lamesch *et al*., 2012) on Phytozome v13 (Goodstein *et al*., 2012). Syntelogs to AtRca (i.e., shared positional homology along chromosomes) were subsequently identified in monocots (*Setaria viridis* Rca: Sevir.8G261433) and asterids (*Mimulus guttatus* Rca: Migut.F00758), which have homology to this sequence using either synteny mapping features on CoGe (Lyons & Freeling, 2008; Lyons *et al*., 2008). Setaria Rca was chosen as an example of a gene that has been previously characterized (Kim et al., 2021) while Mimulus shares comparable synteny to Arabidopsis in the genomic region surrounding Rca enabling identification of syntelogs in the asterids. Working from these three representatives, additional syntelogues were locally identified based on their closest known Rca sister lineage within the Angiosperm phylogeny across other accessible plant genomes on these databases. *Amborella trichopoda* was leveraged as an outgroup for flowering plants in this case study given that this is known to be the one of the earliest diverging lineages for flowering plants (Soltis *et al*., 2011). Additionally, homologues were surveyed, which contained threonine at position 78 phosphorylation site even if they didn’t precisely fit this stringent criterion of synteny using pBLAST on NCBI (https://blast.ncbi.nlm.nih.gov/Blast.cgi) for key agronomic model systems. This ultimately resulted in 217 isoform CDS sequences being identified. Alternative splicing of the same locus in some lineages meant that these CDS sequences represented 153 unique genes from across 91 species and 50 plant families. The longer, Rca-α, sequence was preferentially retained as a taxonomic representative for each alternative splicing gene when available for phylogenetic analysis. The CDS sequence of these genes were visualized in Seaview (Gouy *et al*., 2010) and codon-aligned using MUSCLE (Edgar, 2004). Obvious pseudogenes that were missing large portions of their AAA+ ATPase domain such as the rice sequence (LOC_Os11g47980) were removed prior to phylogenetic analysis of this DNA alignment. A maximum likelihood tree was subsequently assembled using IQTree (Nguyen *et al*., 2015) with the Amborella Rca sequence being specified as an outgroup and 1000 bootstraps being run to generate statistical support for nodes along the tree with the automatic substitution model selection determining a SYM+R8 model was the best-fit according to Bayesian Information Criterion. The resulting treefile was then visualized either in Figtree (http://tree.bio.ed.ac.uk/software/figtree/) or in R using the Phytools and Ape libraries (Paradis & Schliep, 2019; Revell, 2024). Ancestral state reconstructions were performed using the Ape function *ace* particularly for the original position 78 amino acid identity between syntelogous genes, and the manner of α/ß origin (either by alternative splicing or distinct genes) for each plant family (Figure S3).

### Plant growth and stress treatment

Arabidopsis Col-0 and camelina seeds were sterilized in 1 % (v/v) bleach for 5 min, rinsed with 70 % EtOH, and dried on filter paper. The sterilized seeds were sown on MS (Murashige and Skoog basal salt mixture powder, Phyto Technology Laboratories, M524) agar (0.8 %, w/v, Sigma-Aldrich, A1296) with 1 % (w/v) sucrose. The plates were stored at 4 °C for 2 days for cold stratification before transfer to a growth chamber. Seedlings were transferred into pots (5 cm diameter) filled with soil (Berger; PRO-MIX BX All Purpose Growing Mix) before the third true leaf emerged, which was approximately 5 to 7 days after germination (DAG). The other plants (brassica, soybean, green bean, and Medicago) were directly sown in soil after sterilization. All plant species were grown at 22 °C, 16 h light and 8 h dark, 200 µmol m^−2^ s^−1^ PPFD with 2100 K 400W high pressure sodium bulbs (Ceramalux ALTO; C400S51).

### Protein extraction and immunoblotting assay

Total proteins were extracted from three frozen leaf discs (1 cm in diameter) using a Qiagen Tissuelyzer for tissue grinding. The ground tissue was suspended in 0.5 mL of extraction buffer containing 62.5 mM Tris-HCl (pH 6.8), 2% SDS, 10% glycerol, 0.35 M 2-mercaptoethanol, and 0.001% protease inhibitor cocktail (Sigma-Aldrich; P9599). The suspension was heated at 95°C for 5 minutes to denature proteins and inactivate proteases, followed by centrifugation at 13,500 rpm for 30 minutes at 4°C. The supernatant was collected, and 10 µL aliquots were separated on 10% SDS-PAGE gels. Proteins from the gels were transferred electrophoretically to 0.2 µm nitrocellulose membranes (Bio-Rad) and blocked with PBS solution with 5% (w/v) non-fat milk powder (Sigma) for 1h at room temperature. For immunodetection, primary antibodies were diluted in Intercept Blocking Buffer (LI-COR; 927-70001); anti-Rca (Agrisera; AS10 700) at 1:5000 and anti-pT78 (GenScript) at 1:4000, and incubated overnight at 4°C. Modification-specific antibody was generated against the phosphopeptide antigen RGLAYDpT78DDQQDC (Arabidopsis anti-pT78). For each peptide antigen, the Cys residue at the N- or C-terminus was added for coupling to keyhole limpet hemocyanin (KLH). All custom antibodies were produced by GenScript (Piscataway, NJ, USA). Secondary detection used a fluorescent goat anti-rabbit antibody (Invitrogen; A32735) at a 1:10,000 dilution in PBS-T, incubated for 1 h at room temperature, and visualized with a LI-COR Odyssey Infrared Imaging System. All immunoblotting experiments were conducted at least twice to ensure reproducibility of results.

### Protein phosphorylation assay with phosphatase inhibitor

WT Col-0 plants were grown as described above and whole rosettes were sampled and dipped in the protein phosphatase inhibitor buffer: 1% (v/v) solution of phosphatase inhibitor cocktail 2 (Sigma: P5726) and phosphatase inhibitor cocktail 3 (Sigma: P0044), overnight under dark. For dark treatment samples, the protein extraction was done using the overnight incubated samples and for light treatment samples the rosettes were exposed to 150 µmol m^−2^ s^−1^ PPFD light for 2 h and then total protein was extracted.

## RESULTS

### Thr 78 in Rca is common

To study the conservation of Thr78 (Arabidopsis numbering) in Rca, 1893 protein sequences from 479 plants species were assessed. Among these, 203 species retained Thr78 (Table S1). Next, a more stringent synteny informed ortholog phylogeny of Rca-α and Rca-β isoforms was generated across angiosperms to address the questions: at what stage of flowering plant evolution did the putative Thr78 phosphosite emerge, and how well conserved is it among photosynthetic angiosperms? Leveraging maximum likelihood gene family phylogeny ancestral states, Thr78 distribution was estimated for each family, including deeper nodes within the tree such as the branch point of the *Amborellaceae* (Fig. 1). The presence of Thr78 was observed in most plant families (Fig. 1), while among the *Poaceae*, that includes major crop species, the presence of Isoleucine (Ile) was found to dominate at position 78 compared to Threonine (Fig. 1). The difference in the abundance of both amino acids, Thr and Ile at this site is similar at the evolutionary scale, and an estimate of the ancestral state was ambiguous. Interestingly, the a and f3 copies appeared polyphyletic, with multiple origins of the two isoforms arising from alternate splicing or gene duplication across the phylogeny (Fig. S1-S3). Within the Brassicaceae *Arabidopsis thaliana* and its immediate relatives all appear to have a single copy of Rca with alpha and beta forms being the result of alternative splicing of a single transcript.

**Figure 1:**
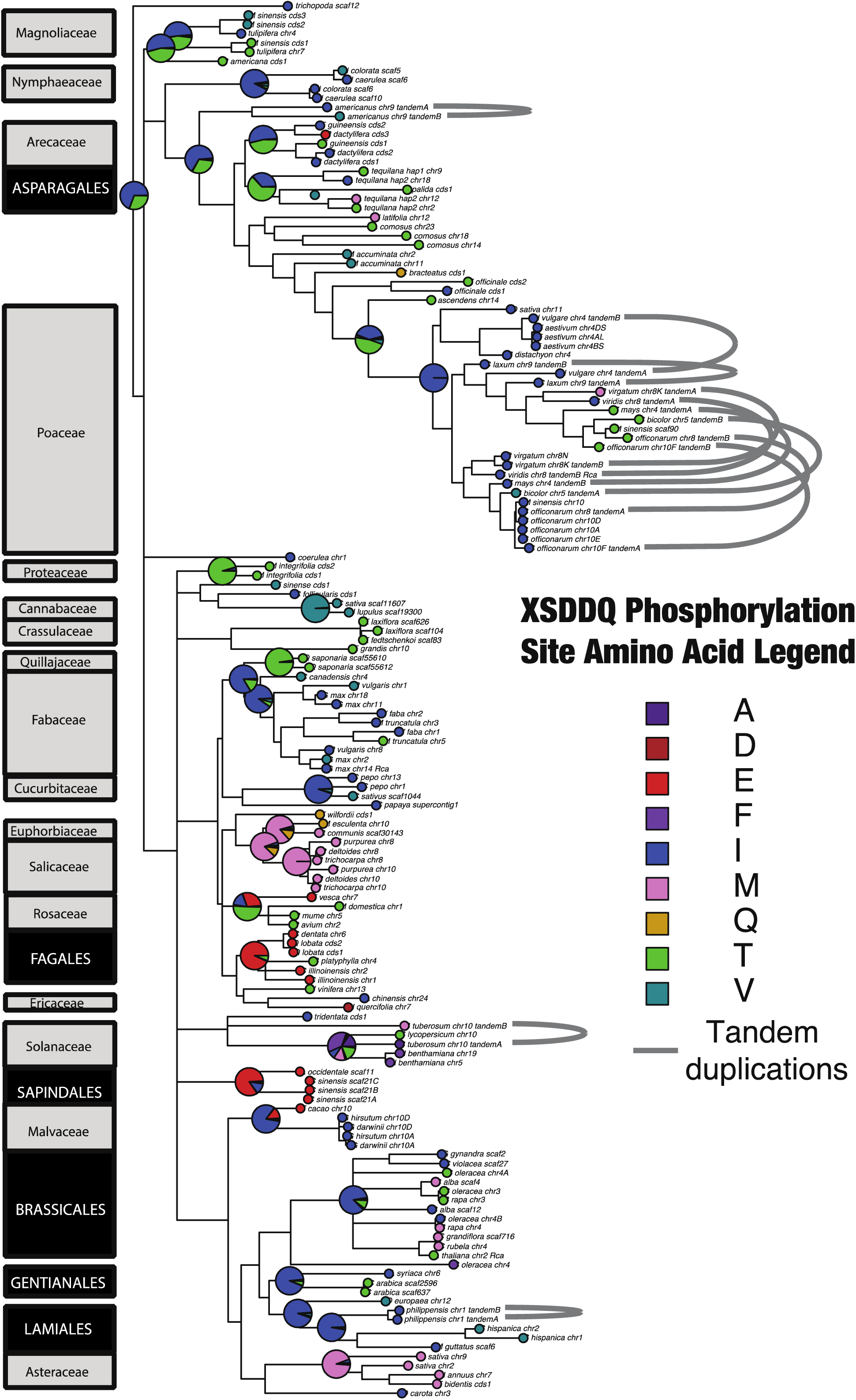
Phylogenetic representation of the amino acid present at the phosphorylation site (peptide position 78) within extant Rca gene models. Ancestral states for families, and when available deeper orders of Angiosperms, are denoted at their corresponding nodes as pie charts. The nodes depicting the presence of Thr78 is colored green and the ones with Ile78 is colored navy blue. Less abundant amino acids at the peptide position 78 are depicted as varying shades of blue and purple. The detailed confirmation of bootstrap values is mentioned in Figure S3.

By contrast in the Fabaceae, *Glycine max* has four gene copies of Rca (two each of a and f3) with the Papillionoid legumes more generally having two copies of Rca. There is no equivalent duplication event in *Cercis canadensis* that would indicate a Fabaceae-wide pattern. Therefore, this pattern of two Papillionoid copies is consistent with the whole-genome doubling (WGD) event preceding the diversification of this subfamily, and by undergoing an additional allopolyploid event, *Glycine max* has two additional copies for each of these ancestral homeologues (Gill, *et al*., 2009; Koenen, *et al*., 2021). Within Poaceae, *Setaria viridis* and the grasses more generally displayed two mirrored clades, which reflect the known taxonomic sorting of this family. The grass family has undergone three WGD events (τ, σ, and ρ), two of which are shared with monocots, with the p WGD unique to the Poaceae (Paterson *et al*., 2004; Tang *et al*., 2010; McKain *et al*., 2016). Although superficially resembling the p WGD event, these grass copies are physically proximal to their sister copies in the adjacent clade suggesting that this may instead be an ancient tandem duplication event.

### Light-dependent dephosphorylation of Thr78 in Rca is unique to Arabidopsis

Previous reports showed that both AtRca isoforms phosphorylate in the dark and are dephosphorylated as plants are exposed to light. To test whether Thr78 phosphorylation status is light-sensitive in species other than Arabidopsis, we screened three C3 plants from the *Brassicaceae* for dark-dependent phosphorylation of Thr78, using a pT78 antibody (Kim et al., 2019) (Fig. 2A). As in Arabidopsis, Rca-α and Rca-β isoforms arise from alternate splicing in *Brassicaceae* and the Thr78 site is conserved in all proteins (Fig. 2B). However, unlike Arabidopsis, Rca in *Camelina sativa* (*Cs*), *Brassica oleraceae* (*Bo*) and *Brassica napus* (*Bn*) phosphorylated in dark but lacked light-induced dephosphorylation (Fig. 3A). These data suggest that although Thr78 is phosphorylated in multiple species, light-dependent regulation may be specific to Arabidopsis.

**Figure 2:**
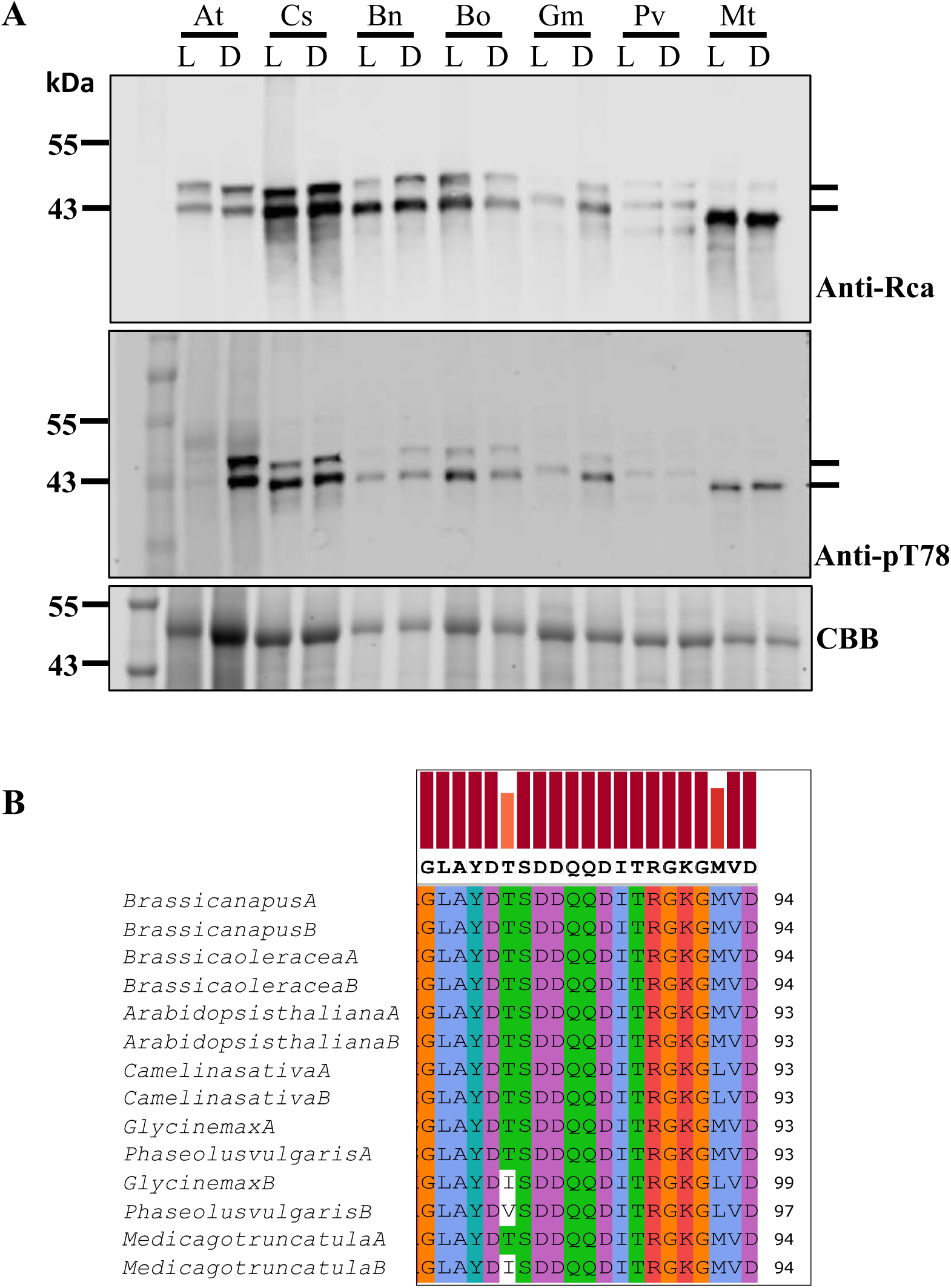
Light dependent dephosphorylation of Thr78 in Rca is specific to Arabidopsis. (A) Immunoblot analysis comparing protein phosphorylation and total abundance of Rca under light (L) and dark (D) conditions (At: Arabidopsis thaliana, Cs: Camelina sativa, Bn: Brassica napus, Bo: Brassica oleraceae, Gm: Glycine max, Pv: Phaseolus vulgaris, Mt: Medicago truncatula). The upper panel shows the total Rca protein extracted from plant samples and the middle panel shows the phosphorylated Rca protein. α and β denote the 2 isoforms of Rca protein. CBB (Coomassie Brilliant Blue) depicts the loading control. (B) Sequence alignment depicting the conservation of Thr78 amino acid in C3 species. The amino acid lacking color codes depicts the change in the Thr78 site to Isoleucine (I) in Glycine max and Medicago truncatula, and Valine (V) in Phaseolus vulgaris. The letter “A” and “B” at the end of the species labelling denotes the Rca-α and Rca-β isoforms. (C) The phylogenetic tree depicts the division of C3 Brassicaceae family and legumes based on Rca isoforms arising from protein splicing in Brassicaceae as compared to different genes in legumes and presence and absence of Thr78 amino acid in the Rca-β isoform. The blue boxes depict the presence of Thr78 amino acid whereas the red depicts the lack of Thr78. The yellow box depicts the experimental confirmation of protein phosphorylation at Thr78 phosphosite.

**Figure 3:**
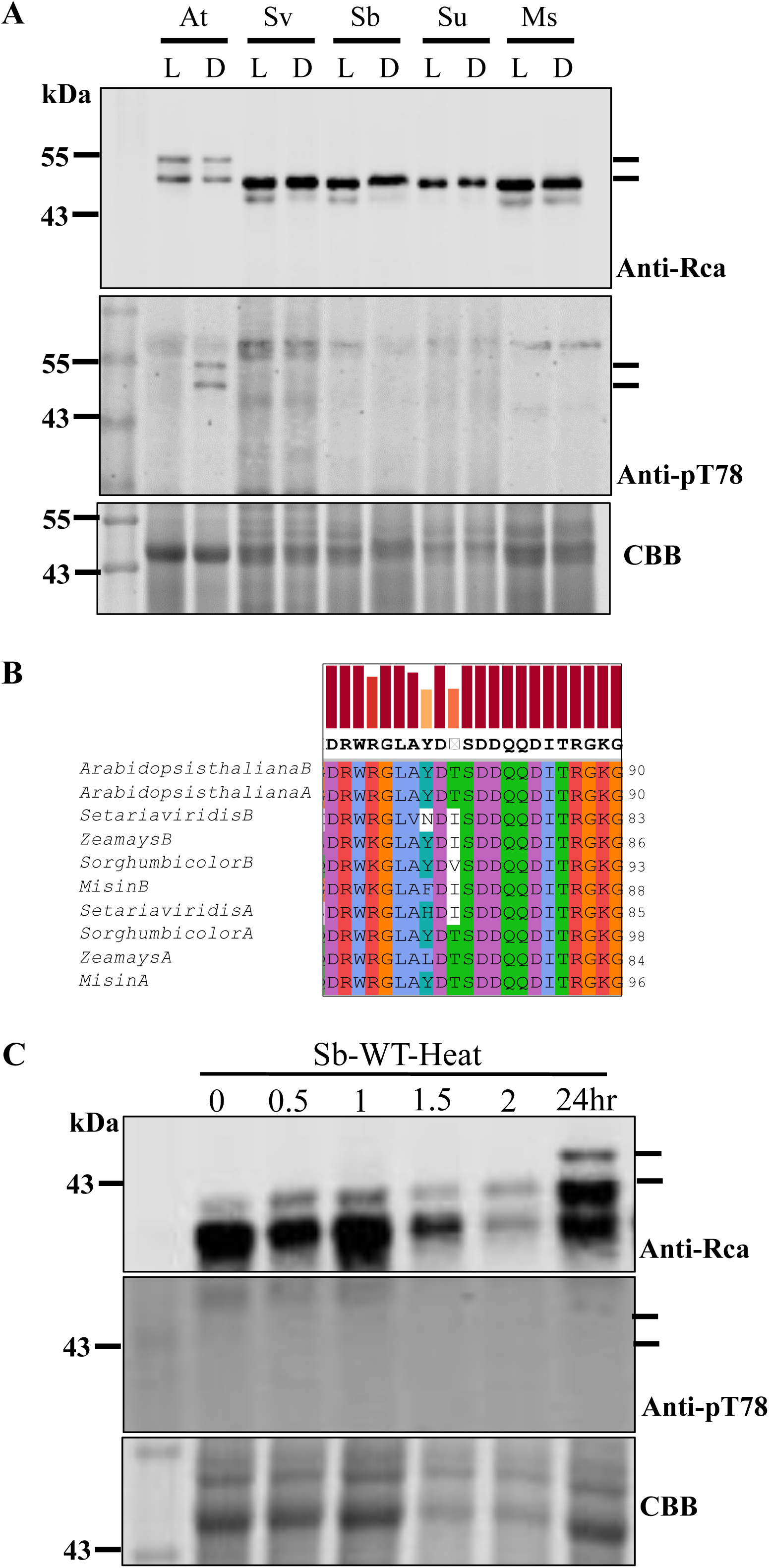
Phosphorylation of Rca is not observed in selected C4 moncots. (A) Protein phosphorylation assay depicting four C4 plant samples in light and dark conditions (At: *Arabidopsis thaliana*, Sv: *Setaria viridis*, Sb: *Sorghum bicolor*, Su: *Saccharum officinarum*, Ms: *Miscanthus*). The upper panel shows the accumulation of Rca protein quantified with Anti-Rca antibody and middle panel shows the phosphorylated Rca protein quantified with Anti-pT78 antibody. α and β denote the 2 isoforms of Rca protein. CBB denotes the loading control. (B) Sequence alignment of motif comparing Arabidopsis and select C4 plants. (C) Immunoblot analysis comparing Rca phosphorylation under heat for 24 h in WT sorghum. The numbers denote the time in hours after the heat stress. CBB denotes the loading control.

To broaden the analysis, we performed the dephosphorylation assay on several legumes: *Glycine max* (*Gm*), *Phaseolus vulgaris* (*Pv*), and *Medicago truncatula* (*Mt*). The Rca isoforms in these species are derived from individual genes of *Rca-α* and *Rca-β*, unlike the situation in the *Brassicaceae*. The Rca-α isoforms retain Thr78, whereas the β isoforms have Ile78 in *Gm* and *Mt,* and Val78 in *Pv* (Fig. 2B). Publicly available data suggest the transcription level of Rca-α is low relative to β in both *Gm* and *Pv,* and moderately expressed in *Mt* (Table S2). The light/dark assay on legume plants showed that both the isoforms were expressed, and phosphorylated in the dark, but as with the Brassicaceae, did not dephosphorylate in the light (Fig. 2A). The reason why the antibody was able to detect phosphorylation of β forms, which lack Thr78, is unclear. It is possible that there is relaxed selectively, but we consider these results with caution.

To study the impact of phosphorylation in C4 grass plants, we analyzed Rca phosphorylation in four species of C4 grass: *Sorghum bicolor* (*Sb*), *Miscanthus* (*Ms*), *Saccharum officinarum* (*Su*), and *Setaria viridis* (*Sv*) (Fig. 3A). Each of these species possess two separate genes for the alternative Rca isoforms, with Rca-α containing Thr78, but with it lacking from Rca-β (Fig. 3B, C). In accordance with previous reports (Kim *et al*., 2021), only the Rca-β isoform could be detected under normal growth conditions (Fig. 3A and C). An unspecific cross-reaction was present in C4 grass samples probed with pT78 antibody, but the detected protein was too large to be Rca-β, subsequently no phosphorylation of Rca-β was detected (Fig. 3A). To test whether a C4 Rca-α isoform could be phosphorylated, sorghum plants were exposed to heat induce Rca-α isoform expression (Fig. 3C). Plants accumulated SbRca-α after 24 h heat treatment, but no bands of the correct size could be detected using the pT78 antibody (Fig. 3C). To rule out the possibility that a lack of detectable Thr78 phosphorylation was due to high protein phosphatase activity, the assay was repeated in the presence of a protein phosphatase inhibitor cocktail (PPIC), but again protein phosphorylation was not observed under any condition (Fig. 4). This result suggests that SbRca-α T78 is not phosphorylated by cpCK2 as it is in Arabidopsis even though the gene for cpCK2 is present in sorghum.

**Figure 4:**
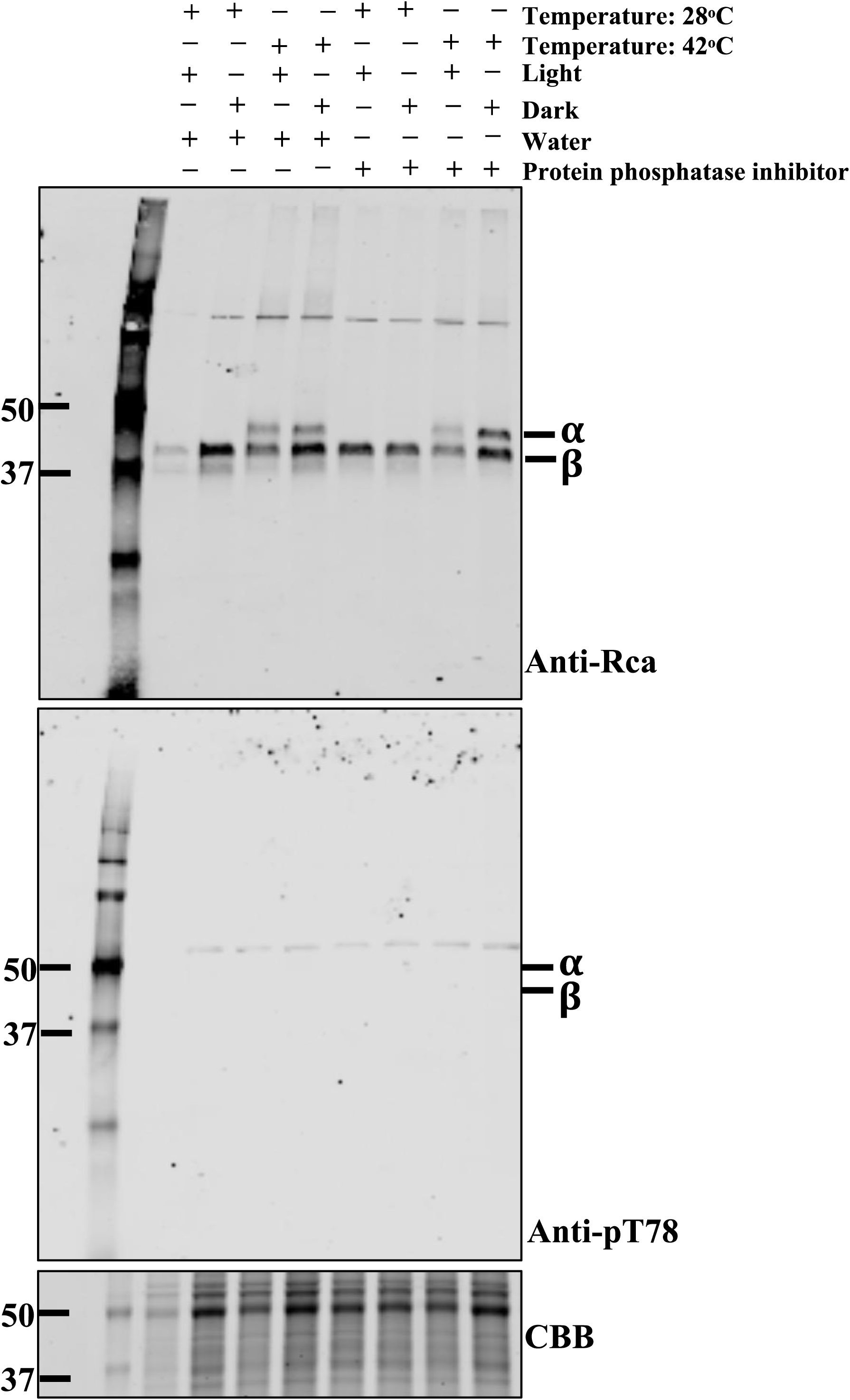
Sorghum lacks phosphorylation of Thr78 with or without PPIC treatment. (A) Immunoblot analysis comparing sorghum WT samples collected after heat stress at 42°C under light and dark treatment. The control samples were collected at 28°C. The + and – sign denotes presence and absence of treatment, respectively. The left panel shows Rca protein accumulation and right panel depicts analysis using anti-pT78 antibody.

## DISCUSSION

Rca is subject to multiple levels of regulation to adjust carbon fixation in response to environmental conditions (Amaral *et al*., 2024), and differences in gene expression, biochemical properties of isoforms, redox status, and post-translational modification of Rca have been topics of investigation (Boex□Fontvieille *et al*., 2014; Kim *et al*., 2016, 2019). Dark dependent phosphorylation of Thr78 is proposed to downregulate Rca activity leading to reduced assimilation (Kim et al. 2019). cpCK2 is described as the major kinase responsible for phosphorylation of AtRca Thr78 based on dramatically reduced phosphorylation in *cpcpk2* T-DNA insertion lines (Kim *et al*., 2016). Substitution of Thr78 with Serine lead to a relative increase in phosphorylation at the site, in Arabidopsis resulted in smaller plants with decreased photosynthesis (Kim *et al*., 2019). Altering a single amino acid, Thr78, was proposed as a means of generating plants with better photosynthesis. However, the prevalence of Thr78 across species, the frequency of post-translational modification and functionality were unclear. To begin to address these questions we expanded previous searches of genomic databases, which assessed the evolution of Rca copy number across the green lineage (Nagarajan & Gill, 2018; Kim *et al*., 2019), to use a synteny based approach. Nagarajan and Gill (2018) placed the emergence of the Rca-α CTE and alternate splicing mechanisms in the charophytes, before the divergence of land plants and there is only one reported example of a species only having evidence for expression of Rca α (Nagarajan & Gill, 2018). Here, despite α and β forms (bearing or lacking a c-terminus respectively) occurring within each model system, these α and β forms appear to represent repeated cases of evolutionary convergence on these two isoforms (Fig S5) through distinct means within each clade: subfunctionalization (Birchler & Yang, 2022), following tandem duplication in the grasses as previously described (Nagarajan & Gill, 2018), WGD in the legumes, or alternative splicing of a single coding sequence into alternative isoforms in Brassicaceae. This may suggest that Rca isoforms can be quite plastic in the way they arise within distinct plant families where α and β forms may not necessarily reflect true homology within Rubisco activase isoforms.

Previous assessment of a smaller number of Rca sequences failed to identify any with Thr78 replaced by Serine, leading to speculation that the configuration with the attendant hyper phosphorylation observed in Arabidopsis has been selected against (Kim *et al*., 2019). In the initial set of Rca sequences identified by BLAST search, we identified gene models from at least seven species (Table S1) which do have Ser78, although these were not considered orthologs in the synteny analysis. Of the 25 species found to contain Rca sequences with Thr78, it remains to be seen how many undergo phosphorylation. Our data are consistent with it occurring in at least the Brassicaceae, while phosphorylation of an equivalent Thr residue has been reported in tomato, suggesting this modification may occur more widely (Marques et al., 2021).

We found no evidence for phosphorylation of Thr78 in monocot species, including Sorghum, Setaria, sugarcane and Miscanthus. The peptide fragment used for antibody generation is highly conserved, with 100% identity between Arabidopsis Rca and Sorghum Rca-α “RGLAYDTSDDQQDC” and should therefore have been able to detect phosphorylation if present under the conditions tested. There are several potential explanations for this finding, including an absence of the kinase, differences in localization or expression profiles of the kinase, or regulation of the kinase. In Arabidopsis, Rca is phosphorylated by cpCK2 (Kim et al., 2016). A homologous gene has been characterized in rice (LOC_Os03g55490) (Lu et al., 2015) and syntenic sequences can be found in C4 grasses including Sorghum (Sobic.001G07970) (Carvalho et al., 2020). As Sorghum operates two-celled C4 photosynthesis for an interaction to occur between Rca and cpCK2 both must be expressed in the same cell type. Transcriptomics suggests that cpCK2 is expressed in both mesophyll and bundle sheath cells in maize (Chang et al., 2012), Setaria (John et al., 2014) and Sorghum (Döring et al., 2016), albeit with ∼2-fold mesophyll enrichment in maize and Setaria whereas Rca is localized in the bundle sheath. Even so, it appears that the orthologous cpCK2 is present in the same compartment as Rca (i.e., the bundle sheath cell chloroplast). However, in the monocot species tested Rca-β, which lacks Thr78, is the predominate isoform under normal conditions and Rca-α, which possess Thr78, is induced during stress (Shivhare & Mueller-Cajar, 2017; Scafaro et al., 2019; Kim et al., 2020, 2021). Whether C4 monocot cpCK2 retains the ability to phosphorylate Thr78 in Rca-α, or if the bundle sheath stromal environment is permissive for phosphorylation remains unclear (Kim et al., 2016). CK2 exists as a multigene family, composed of catalytic α subunits and regulatory beta subunits (Salinas et al., 2006; Schweer et al., 2010), which form hetero-tetramers that can differ in substrate specificity (Domańska et al., 2005). Intriguingly there appears to be a lack of regulatory β subunits in the chloroplast (Salinas et al., 2006), so differences in regulatory subunits impacting specificity appears unlikely, although multiple chloroplast CK2 containing complexes have been detected (Baginsky et al., 1997) suggesting the protein does not act alone. However, further research is required to characterize the regulation of monocot cpCK2 and whether it can phosphorylate Rca.

Conservation of phosphorylation of Thr78 in some C3 plants is not evidence of a conserved regulatory role. Indeed, in the species examined, strong evidence for light-dependent dephosphorylation of Thr78 was only found for Arabidopsis, with phosphorylation of Rca detected in both the light and dark in other C3 species (Fig. 3). These data may suggest a limited scope for phosphorylation as a means of regulation, with most species depending on other mechanisms, such as changes in redox state, catalytic properties or expression. It may be that alternate post-translational modifications of Rca have greater importance in other species and several different ones have been described (Amaral et al., 2024).

Taken together our results suggest that Thr78 can be found in Rca sequences across the green lineage, although the prevalence varies, and degree to which it undergoes phosphorylation is inconsistent. It is likely that regulation of Rca can occur through multiple mechanisms, including redox, sensitivity to ADP and gene expression and the relative importance of each varies.

## Acknowledgments

We would like to thank Toby Kellogg and Shawn Abrahams for providing feedback on the phylogenic analyses. This work was funded by the DOE Center for Advanced Bioenergy and Bioproducts Innovation (U.S. Department of Energy, Office of Science, Biological and Environmental Research Program under Award Number DE-SC0018420). Any opinions, findings, and conclusions or recommendations expressed in this publication are those of the author(s) and do not necessarily reflect the views of the U.S. Department of Energy.

## Authors contributions

Conceptualization (SK, NB, SJB, DO), Investigation (NB, SC, MS, JH, SK), Formal Analysis (NB), Project administration (NB, SJB, DO), Supervision (AL, SJB, DO), Visualization (NB), Writing - original draft (NB, JH, SC, DO, SJB), Writing – review and editing (NB, JH, DO, AL, SJB).

