## Supplementary figures and images for "Phylogenetic diversity of light dependent phosphorylation of Thr78 in Rca"

### Fig. S1

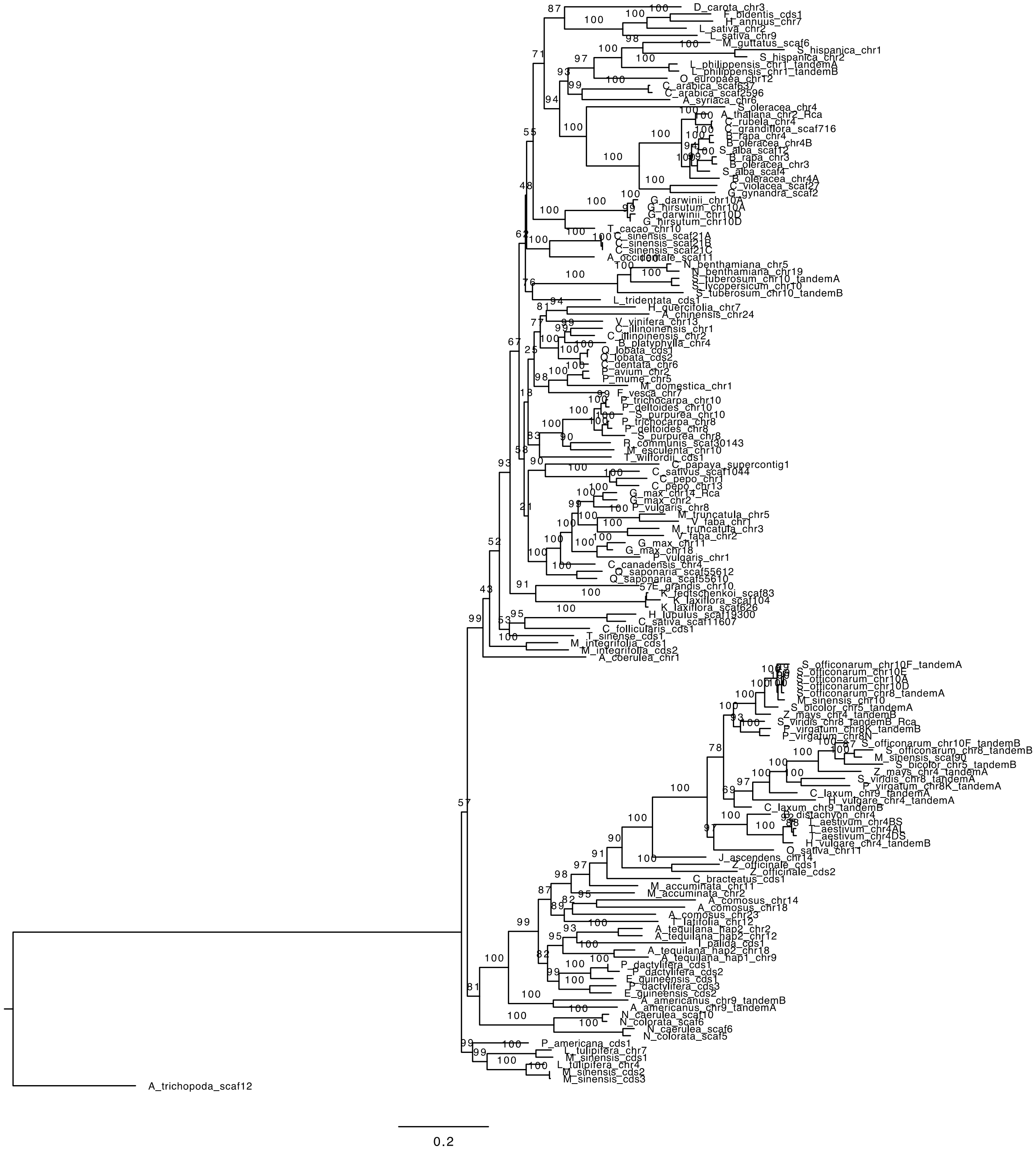

### Fig. S2

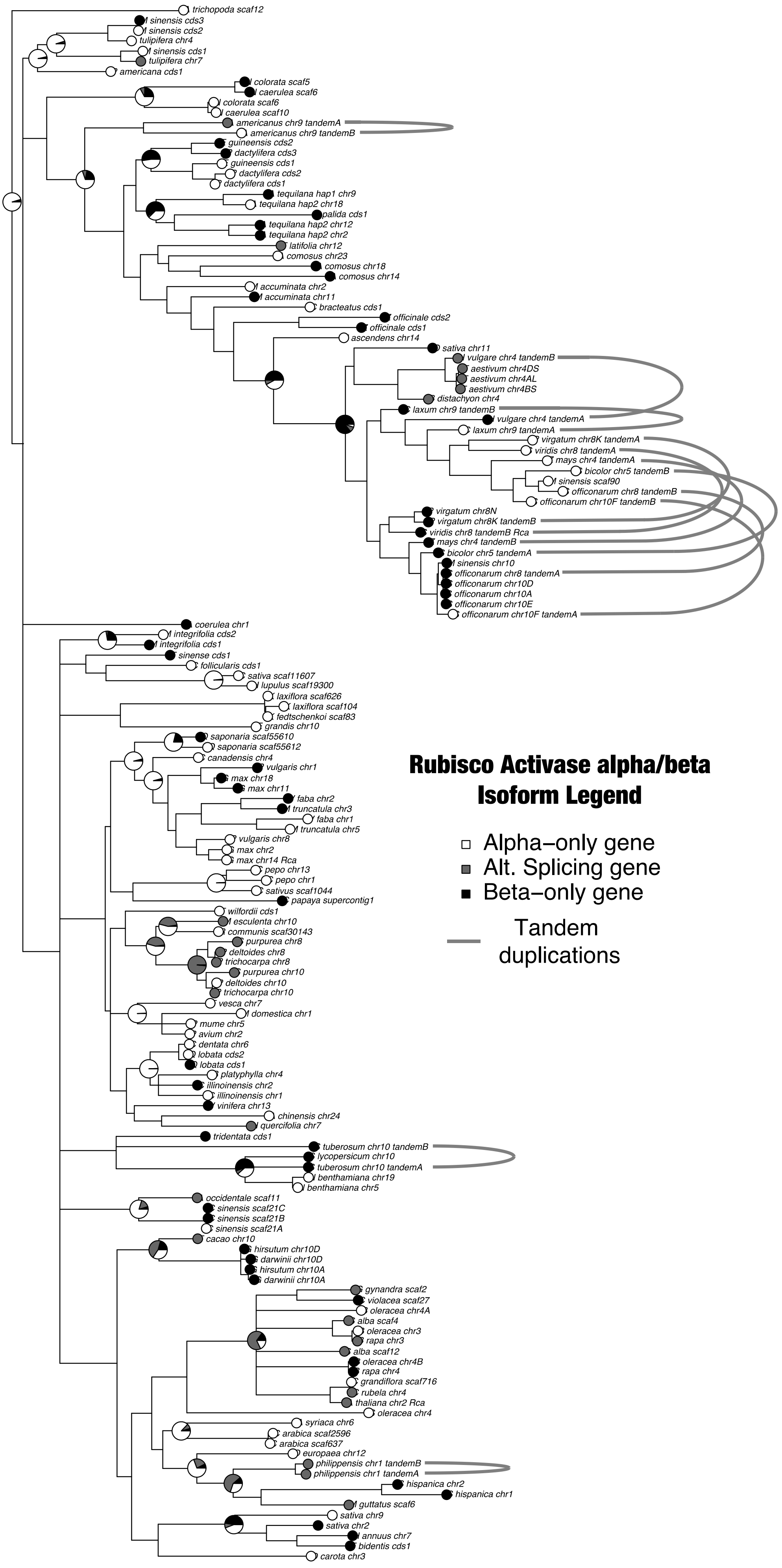

### Fig. S3

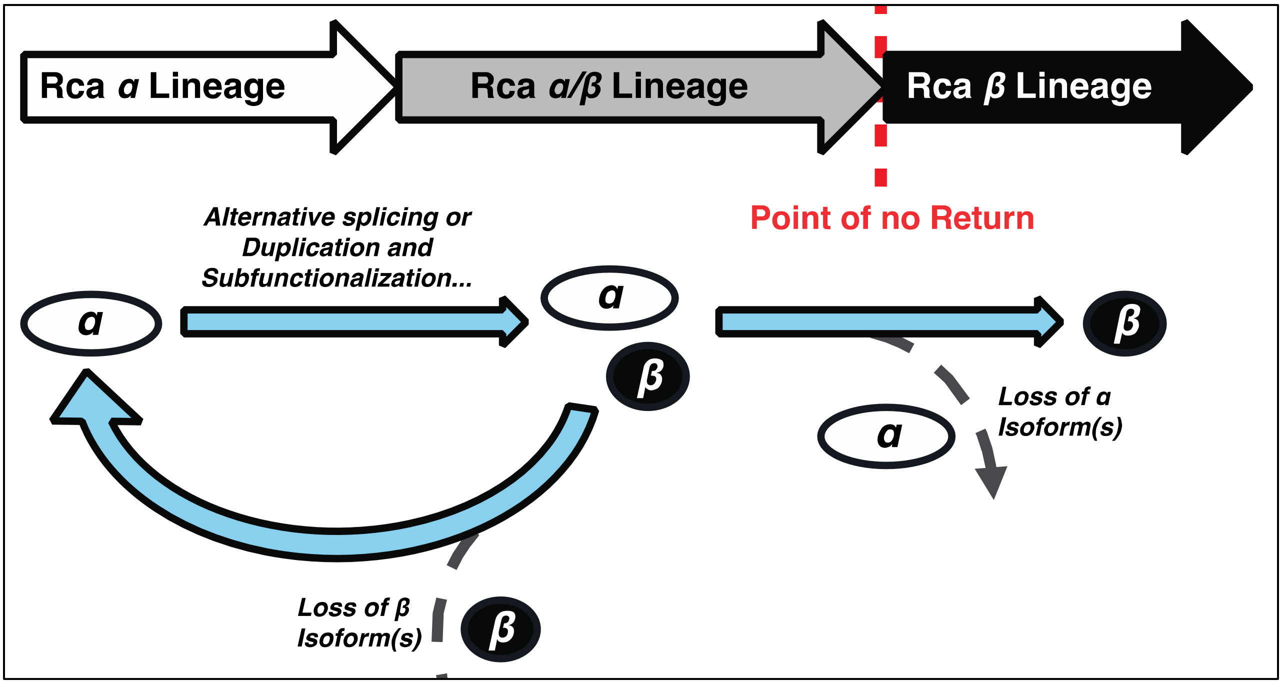
