## Supplementary material for "Phylogenetic diversity of light dependent phosphorylation of Thr78 in Rca": Table S2

| | Gene ID | Protein Isoform | Expression | Amino acid |
|---|---|---|---|---|
| Glycine max | Glyma.02G249600 | a | 312.814 | Val (V) |
| | Glyma.11G221000 | b | 663.672 | Iso (I) |
| | Glyma.14G067000 | a | 193.648 | Iso (I) |
| | Glyma.18G036400 | b | 284.853 | Iso (I) |
| Medicago truncatula | Medtr3g068030 | b | 7725.380 | Iso (I) |
| | Medtr5g080450 | a | 243.074 | Thr (T) |
| Phaseolus vulgaris | Phvul.001G234300 | b | 6766.310 | Val (V) |
| | Phvul.008G192400 | a | 1421.350 | Iso (I) |
